# An outbreak of *Pseudomonas aeruginosa* infection linked to a “Black Friday” piercing event

**DOI:** 10.1101/149914

**Authors:** Peter MacPherson, Katherine Valentine, Victoria Chadderton, Evdokia Dardamissis, Iain Doige, Andrew Fox, Sam Ghebrehewet, Tom Hampton, Ken Mutton, Claire Sherratt, Catherine M McCann

## Abstract

**Background:** Outbreaks linked to cosmetic piercing are rare, but can cause significant illness. We report the investigation and management of a point-source outbreak that occurred during a “Black Friday” event in North West England.

**Methods:** Outbreak investigation was led by Public Health England, and included active case finding among individuals pierced at a piercing premises between 25/11/2016 (“Black Friday”) and 7/12/2016. Detailed epidemiological, environmental (including inspection and sampling), and microbiological investigation was undertaken.

**Results:** During the “Black Friday” event (25/11/2016), 45 people were pierced (13 by a newly-appointed practitioner). Eleven cases were identified (7 microbiologically-confirmed, 2 probable, and 2 possible). All cases had clinical signs of infection around piercing sites, and five required surgical intervention, with varying degrees of post-operative disfigurement. All confirmed and probable cases had a “scaffold piercing” placed with a guide bar by the newly-appointed practitioner. *Pseudomonas aeruginosa,* indistinguishable at nine-locus variable-number tandem repeat loci, was isolated from four of the confirmed cases, and from pre- and post-flush samples from five separate water taps (three sinks) in the premises. Water samples taken after remedial plumbing work confirmed elimination of *Pseudomonas* contamination.

**Conclusions:** Although high levels of *Pseudomonas* water contamination and some poor infection control procedures were identified, infection appeared to require additional exposure to an inexperienced practitioner, and the more invasive scaffold piercing. A proactive collaborative approach between piercers and health and environmental officials is required to reduce outbreak risk, particularly when unusually large events are planned.

## Introduction

Scaffold ear piercing (sometimes known as “bar”, “industrial”, or “construction” piercing) involves the placement of two pierced holes in the upper ear cartilage (one in the forward helix position, and one in the posterior helix), through which a metal jewellery bar with securing ends is threaded, sometimes using a guide bar^1^. Although scaffold piercing is reported to be more painful than conventional ear piercing (as it involves the helix rather than the pinna) and may take longer to heal^2^^-^^4^, we are unaware of previous reports of rates of infection or outbreaks associated with this relatively new cosmetic procedure^5^.

A small number of case reports and point source outbreaks of *Pseudomonas aeruginosa* infection linked to conventional ear piercing have previously been reported^6^^-^^8^. In 2000 in Oregon, USA, cartilage infection was confirmed in seven cases and suspected in a further 18 individuals, with risk of infection strongly associated with piercing with an open gun^7^. In England during 2016, a national outbreak of *Pseudomonas* infection linked to a contaminated aftercare cleaning solution resulted in a large number of cases^9^.

During early December 2016, a local Ear, Nose and Throat (ENT) surgeon working in a District General Hospital notified Public Health England (PHE) North West that, over the course of two days, he had operated upon the ear cartilage of two teenage girls who had both developed infection after undergoing scaffold piercing on the same day and at the same premises. Here we report subsequent investigations and public health control measures implemented to manage the outbreak.

## Methods

In England, piercing premises are regulated by local authorities and they must be registered and undergo a satisfactory hygiene inspection by Environmental Health Officers^10^. All registered persons and premises must comply with the Local Authority Byelaws. There are also recommendations for infection control, set out in guidance jointly produced by the Chartered Institute of Environmental Health, Public Health England, the Health and Safety Laboratory, and the Tattoo and Piercing Industry Union^11^, which operators are encouraged to follow.

The piercing premise involved in this outbreak was located in North West England, was licensed, and had previously undergone satisfactory inspection in June 2015. There were two piercing practitioners: a more experienced owner, and a newly-employed practitioner.

Using social media, the premises advertised a special “Black Friday” event for the 25^th^ November 2016, where all piercings were reduced in price. Following initial reports of infection made by the surgeon, an Environmental Health Officer visited the premises on 8th December 2016, and found that the owner had voluntarily suspended piercing following reports of illness he had received through social media platforms. The Environmental Health Officer and a Community Infection Control Nurse subsequently conducted a detailed environmental and hygiene inspection on 13^th^ December 2016.

An outbreak control team was convened, and comprised of public health specialists, microbiologists, environmental health officers, community infection control nurses, and clinicians. We undertook active case finding by systematically contacting (by telephone, or by letter if unsuccessful) all individuals who, according to the premises’ records, had been pierced between 25/11/2017 and 7/12/2016 (when the owner had voluntarily suspended piercing). Contacted individuals were asked to complete an epidemiological questionnaire and were referred for clinical assessment as appropriate. We also sent letters to all individuals who had been pierced during this period, alerting them to the symptoms of infection and providing a link to complete an online version of the epidemiological questionnaire.

Confirmed infection was defined by isolation of *Pseudomonas aeruginosa* by culture from the pierced site in an individual pierced at the premises between 25/11/2016 and 7/12/2016. Probable infection was defined as an individual pierced at the premises between 25/11/2016 and 7/12/2016 who had clinical symptoms or signs of infection at the piercing site (redness, hotness, tenderness, swelling or discharge), and who had been diagnosed or treated for infection by a clinician. Possible infection was defined as an individual pierced at the premises between 25/11/2016 and 7/12/2016 and who reported symptoms or signs of infection at the piercing site, but who had not been diagnosed or treated by a clinician.

Culture of specimens taken from cases was performed at local National Health Service Microbiology laboratories, and, where possible, isolates of *Pseudomonas aeruginosa* were sent to the Public Health England National Antimicrobial Resistance and Healthcare Associated Infections Reference Laboratory, Colindale, London, England for molecular subtyping by nine-locus variable-number tandem repeat (VNTR) amplification.

We inspected the premises’ sterilising autoclave records to confirm functionality and timing of use, and collected environmental specimens from potential sources inside the premises, including: water samples from hot and cold taps (pre- and post-flush); from disinfectant spray bottles (liquid samples and swabs from spray nozzles); from piercing equipment (swabs from scaffold guide bars); and from the autoclave (solution sample). Additional water samples were taken from the mains water supply and surrounding properties by the water utility company. Environmental samples were cultured for *Pseudomonas* and subjected to nine loci VNTR amplification at the PHE Food and Water Laboratory, York, England.

Using epidemiological data, we plotted an epidemic curve, and calculated attack rates stratified by the type of piercing and practitioner who carried out the procedure.

## Results

### Active case finding and epidemiology

In total, 45 individuals were pierced at the premises during the 25^th^ November 2016 “Black Friday” event (44 female, one male). Mean age was 20 years (range 14-47 years). A further seven individuals were pierced between the 26^th^ November and the 7^th^ December 2016 inclusive, when piercing was suspended.

Of the 45 individuals pierced during the Black Friday event, 13 (29%) were pierced by the newly-appointed practitioner, and 32 (71%) by the more experienced practitioner. Twelve of the thirteen individuals pierced by the newly-appointed practitioner had scaffold piercings placed in the ear by needles, with metal guide bars used to draw the jewellery bar through the pierced holes. The remaining individual had a belly-button piercing placed using the same guide bar technique. One of the 32 individuals pierced by the more experienced practitioner had a scaffold piercing of the ear, with jewellery placed with the use of a guide bar.

None of the seven individuals pierced after the Black Friday event (two pierced by newly-appointed practitioner, and five by owner) had scaffold piercings.

In total 11 cases of infection were identified: seven microbiologically-confirmed; two probable cases; and two possible. Ten of eleven (91%) of cases were female, with ages ranging between 14 years and 36 years. Dates of onset of symptoms ranged from 26^th^ November 2016 to 2^nd^ December 2016, with the epidemic curve indicating a likely point-source outbreak (Figure 1).

**Figure 1:**
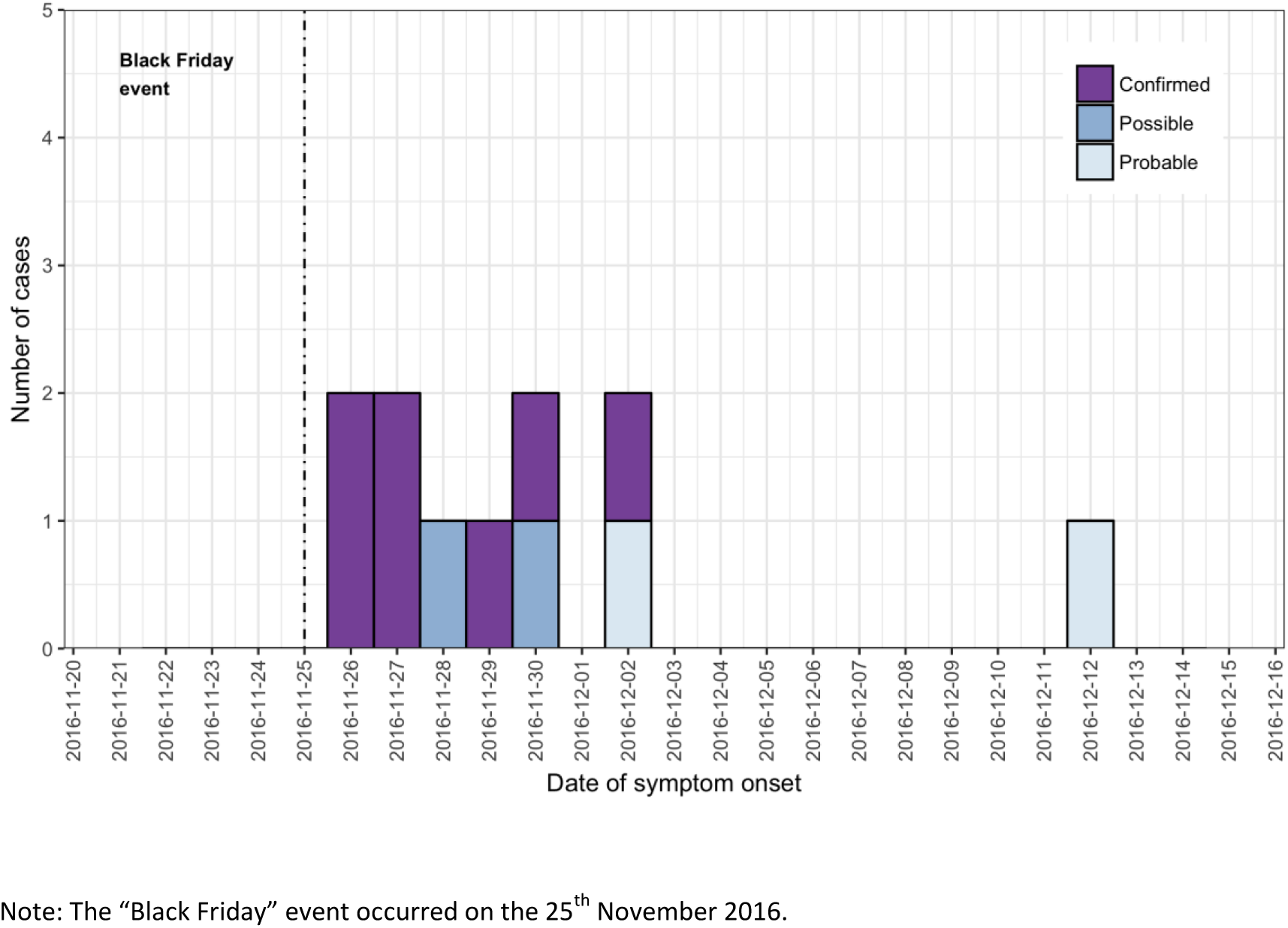
Epidemic curve.

All cases were pierced during the “Black Friday” event by the newly-appointed practitioner, and had scaffold piercing with jewellery placed using guide bars, giving an attack rate of (85%, 11/13) for individuals exposed to the newly-appointed practitioner during the “Black Friday” event, and an attack rate of 0% (0/32) for individuals pierced by the more experienced practitioner during the “Black Friday” event (overall “Black Friday” premises attack rate: 24%, 11/45) – Figure 2.

**Figure 2:**
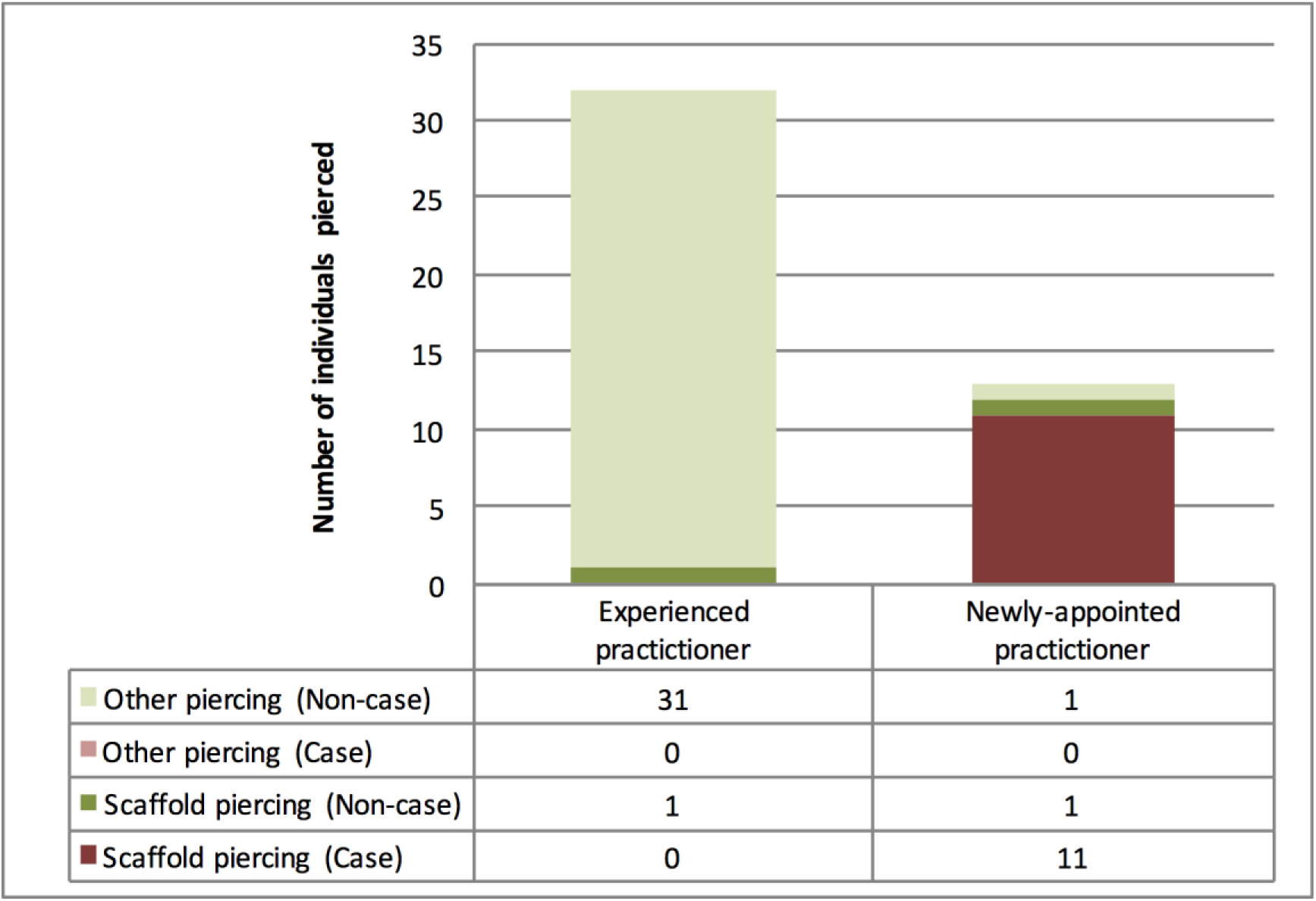
Attack rates by practitioner and procedure among individuals pierced during “Black Friday” event.

Five cases were admitted to hospital and required surgical intervention and antibiotic treatment, and a further four were treated as outpatients with antibiotics only. At least one individual who underwent surgery was left with residual scarring.

### Microbiological and environmental investigation

VNTR typing was completed on 4/7 *Pseudomonas aeruginosa* isolates from microbiologically-confirmed cases, with all isolates being clade C type: 11.6.2.2.1.3.7.2.11, and indistinguishable between cases at the last most discriminatory loci.

Inspection of plumbing and fixtures identified plastic pipes supplying the sinks in the premises. Ten water samples were taken from five taps at three sinks (two pre-flush and two post-flush). Overall, eight water samples grew *Pseudomonas aeruginosa*, with five having very high levels of colony-forming units per millilitre (Table 1), indicating contamination in the plumbing system supplying the washbasin taps. VNTR subtyping of isolates from water samples matched samples taken from cases at the last most discriminatory loci. Water samples from the main supply into the property and from the adjacent property did not grow *Pseudomonas spp.* Samples taken from the autoclave, disinfectant spray bottles, and piercing equipment (including used guide bars) were also negative.

**Table 1:** Isolation of *Pseudomonas aeruginosa from water samples*

| Sink | Tap | *Pseudomonas aeruginosa* concentration in water |  |
| --- | --- | --- | --- |
|  |  | Pre-flush sample | Post-flush sample |
| 1 (Treatment room) | Cold water tap | 51 cfu / 100mls | ND |
|  | Hot water tap | >100 cfu / 100mls | 8 cfu / 100mls |
| 2 (Cleaning room) | Cold water tap | >100 cfu / 100mls | >100 cfu / 100mls |
|  | Hot water tap | ND | 11 cfu / 100mls |
| 3 (Toilet) | Mixer tap | 1 cfu / 100mls | >100 cfu / 100mls |

ND: Not detected. CFU: colony forming units

Inspection of the premises by infection control nurses identified hygiene issues including: lack of a dedicated handwashing sink; use of reusable cotton towels for hand-drying; inadequate solutions used for environmental cleaning and for disinfection of invasive guide bars; and inadequate hand decontamination.

### Control measures

A qualified plumber replaced all plastic pipes with copper piping, as well as washbasin taps on 6/2/2017. No growth of *Pseudomonas spp.* was obtained from water samples taken from all taps after the completion of this remedial work.

Written infection control and hygiene advice was provided to the owner. Following re-inspection by the Environmental Health Officer and confirmation that all issues had been addressed the premises was permitted to recommence piercing activities.

## Discussion

We report on a large outbreak of *Pseudomonas aeruginosa* infection linked to a relatively new type of cosmetic piercing (scaffold piercing) conducted during a busy “Black Friday” event. Based on molecular subtyping of isolates from cases and from water samples taken from taps in the affected premises, the source of the outbreak was most likely to be contaminated plastic plumbing pipes. All cases occurred among clients who had scaffold-piercing done by the newly-appointed trainee, whilst no individuals pierced by the more experienced practitioner or who did not have scaffold piercing, was infected. We therefore hypothesise that whilst there were infection control issues in the premises and the water supply was contaminated with *Pseudomonas aeruginosa*, development of infection may have required additional exposure to the more invasive scaffold procedure conducted by an inexperienced practitioner.

*Pseudomonas aeruginosa* is ubiquitous in the environment (particularly in low nutrient and oligotrophic environments) and can colonise and form biofilms in plumbing fixtures such as in taps, shower head, and pipes^12^. Contamination of water supplies is a well-recognised source of infection, particularly among immunosuppressed individuals^12^^,^^13^. In this outbreak, isolation of a common environmental strain of *Pseudomonas aeruginosa* from multiple taps and sinks and in high concentrations from both pre- and post-flush samples strongly indicates that the plastic pipes more distal from the sinks were contaminated. It is possible that the unusually high numbers of clients pierced during the Black Friday event would have required greater water usage, and may have resulted in a *Pseudomonas-*contaminated biofilm being dislodged from the plastic pipes, contaminating piercers’ hands or equipment during washing. Infection risk may have been increased as handwashing and decontamination procedures were found to be suboptimal.

Although a small number of previous case reports and outbreaks of *Pseudomonas aeruginosa* linked to cosmetic piercing activities have been reported^2^^,^^6^^,^^7^^,^^9^^,^^14^, we are unaware of outbreaks associated with this new, more invasive type of piercing, or with a large venue-based event such as reported here. Our investigations demonstrate that piercers should work proactively and closely with environmental health officers, infection control nurses and with licensing bodies when planning unusually large events. If such a risk assessment had been conducted in this case, it might have been possible to reduce the risk of infection posed by suboptimal infection control procedures, and from the less-experienced piercer undertaking the more invasive scaffold piercings using guide bars. Molecular subtyping was critical to rapidly identify the source of this outbreak, and to effectively direct control efforts.

There were some limitations to this investigation. Molecular subtyping was successfully completed for only 4/7 isolates; other subtypes may have been identified from non-typed isolates. Probable and possible cases may be attributable to infection with organisms other than *Pseudomonas aeruginosa.* Although the epidemic curve indicates a point-source outbreak, and the rapidity of onset of symptoms suggests delivery of a high bacterial load to individuals during piercing, it is possible that some cases with longer incubation periods resulted from person-to-person transmission, for example through close contact or sharing of cleaning materials.

In summary, the combination of water contamination with *Pseudomonas aeruginosa*, an unusually high number of clients during a Black Friday event, piercing by an inexperienced individual, and the more invasive scaffold piercing resulted in a large outbreak with a very high attack rate. Piercers should proactively work with environmental health officers and infection control specialists when planning large-scale events.

## Acknowledgements

We gratefully acknowledge support from: Cheshire and Merseyside Health Protection Team, PHE North West; PHE Field Epidemiology Service, North West; and Wirral Council.

## Funding

There were no sources of financial support for this study.

## Notes

***Conflict of Interest***: We confirm that all authors have no commercial or other associations that may pose a conflict of interest.

